# PARS: an automated, open-source pipeline for subject-specific finite element head modelling from MRI

**DOI:** 10.64898/2026.07.05.736584

**Authors:** Vahid Darvishi, Emily Yik Kwan Chan, Harry Duckworth, Thomas D. Parker, David J Sharp, Mazdak Ghajari

**Author notes:** Corresponding author: Vahid Darvishi < >.

## Abstract

Converting medical images into anatomically detailed, subject-specific finite element (FE) models is a long-standing bottleneck in brain computational modelling. These models are used to predict brain tissue deformation, e.g. in traumatic brain injury, particle diffusion in brain drug delivery, and other biophysical phenomena across neurological disorders. However, existing model creation workflows depend on manual image segmentation, proprietary meshing software, and labour-intensive repair of meningeal and interface structures, limiting reproducibility and cohort analysis. Here we present PARS, a fully automated, open-source pipeline that converts a T1-weighted MRI scan into a simulation-ready FE head model. PARS combines anatomical parcellation with tissue maps and uses iterative neighbourhood-based reclassification, yielding a gap-free whole-head label volume. The volume is directly converted into a hexahedral mesh, augmented with algorithmically reconstructed falx, tentorium, pia and dura mater, and refined by Laplacian smoothing under a node-locking scheme that controls element quality and the explicit-solver stable timestep. We evaluated PARS on 23 subjects spanning cranial volumes of 832–1,329 cm³, at 1.0, 1.5 and 2.0 mm MRI resolutions. At 1 mm, meshes achieved a median Scaled Jacobian of 0.976±0.012, and total intracranial volume error of 0.54±0.19%; quality remained high at 1.5 mm (SJ: 0.933±0.018) and 2 mm (SJ: 0.921±0.016). Model creation runtime ranged from 9 to 38 minutes per subject. Models generated by PARS have been validated against cadaveric brain displacement data and demonstrated utility across traumatic brain injury and normal pressure hydrocephalus research. PARS provides an open-access, reproducible resource that substantially lowers the barriers to subject-specific brain modelling.

## 1 Introduction

Neurological disorders are a leading cause of death and disability worldwide, with prevalence increasing steadily over recent decades partly due to the rise in the aging population (Feigin et al., 2020). In several of these disorders, including traumatic brain injury (TBI), idiopathic normal pressure hydrocephalus (iNPH), intracranial bleeding, and brain tumour growth, mechanical loading directly contributes to tissue damage. Understanding the relationship between tissue mechanical loading and damage can help us improve prevention, diagnosis and treatment. However, it is nearly impossible to measure tissue loading in the living human brain. Tagged MRI has been used for this purpose, but it is limited to controlled low severity head impacts (Bayly et al., 2021; Lu et al., 2026). In vitro and animal studies have been used to explore links between mechanical loading and brain damage (Browne et al., 2011; Donat et al., 2021; Fournier et al., 2014; Li et al., 2019; Nakadate et al., 2017), but measuring tissue loading remains a challenge and translating the findings to human is limited by anatomical and biological differences.

Finite element (FE) modelling is a powerful method for estimating brain tissue deformation in different neurological conditions. It has been applied most extensively to studying TBI biomechanics (Duckworth et al., 2022; Ghajari et al., 2017; Ji et al., 2014; Kleiven and Von Holst, 2002; Mao et al., 2013; Zimmerman et al., 2023). By incorporating a description of brain anatomy, material properties of different tissues and their interface, and boundary conditions, FE models can predict the brain tissue’s biomechanical response to complex loading conditions at high spatial and temporal resolution (Ghajari et al., 2017; Li et al., 2021; Miller et al., 2016; Zhao and Ji, 2020). However, their adoption has been limited by poor accessibility to FE brain models and reproducibility of models in different labs. Most published models and their codebases are unavailable, and their mesh generation workflows often rely on proprietary software (Ji et al., 2022; Lyu et al., 2022; Ratajczak et al., 2019; Sbriglio et al., 2025). These barriers limit independent validation, reuse and broader deployment. Without a transparent, reproducible, and fully open-source pipeline, the translation of subject-specific brain FE modelling from academic curiosity to clinical utility remains unachievable.

The prediction accuracy of brain models depends on their anatomical fidelity (Liu et al., 2022). Most brain FE meshes are generated using surface-based methods, in which segmented tissues are first represented with smooth surfaces and then discretised using hexahedral or tetrahedral elements (Ji et al., 2014; Kleiven and Von Holst, 2002; Mao et al., 2013). These methods often simplify or omit anatomical features, such as gyri and sulci (Miller et al., 2016), despite evidence that these features produce loading patterns that overlap with pathology distribution (Ghajari et al., 2017; Ho and Kleiven, 2009; Kim et al., 2015; Zimmerman et al., 2023). The surface-based methods also require substantial manual effort, which limits their scalability (Chen and Ostoja-Starzewski, 2010).

Voxel-based meshing offers a more anatomically faithful alternative by converting MRI voxels directly into hexahedral FE meshes, preserving gyral and sulcal geometry without surface reconstruction. The main limitation is stair-step artifacts at tissue interfaces, which can produce unrealistic stress concentrations but can be substantially reduced by mesh smoothing algorithms (Chen and Ostoja-Starzewski, 2010; Zhou et al., 2025). However, existing voxel-based pipelines often require manual segmentation and manual construction of key intracranial structures, such as the falx and tentorium. This is a bottleneck that limits scalability and likely explains why these features are omitted in several published brain models (Chen and Ostoja-Starzewski, 2010; Miller et al., 2016; Zhou et al., 2025).

Here we present PARS (Pipeline for Automated Reconstruction of Subject-specific head models), a fully automated, open-source pipeline for generating anatomically detailed, subject-specific FE head models directly from standard T1-weighted MRI scans. This pipeline integrates: i) robust tissue segmentation, ii) voxel-based mesh generation, smoothing and contact repairing, iii) generation of the falx, tentorium, pia mater and dura mater, and iv) control over mesh quality and stable time step of explicit-integration solvers. We describe the key components of the pipeline. We then use it to automatically generate head FE models for a diverse range of subjects and report outputs, including mesh quality metrics, stable timestep and model creation time. By providing this open-source pipeline, we aim to advance transparent, reproducible and scalable FE modelling of the human brain for studying various neurological conditions.

## 2 Materials and methods

### 2.1 Overview of the pipeline

The model generation workflow of PARS consists of several stages, shown in Fig. 1 and explained in the following sections. These include: reading the images generated by FreeSurfer’s recon-all, image pre-processing, hybrid segmentation, voxel-to-element mesh generation, algorithmic reconstruction of meningeal structures, mesh smoothing and contact repair.

**Fig. 1.**
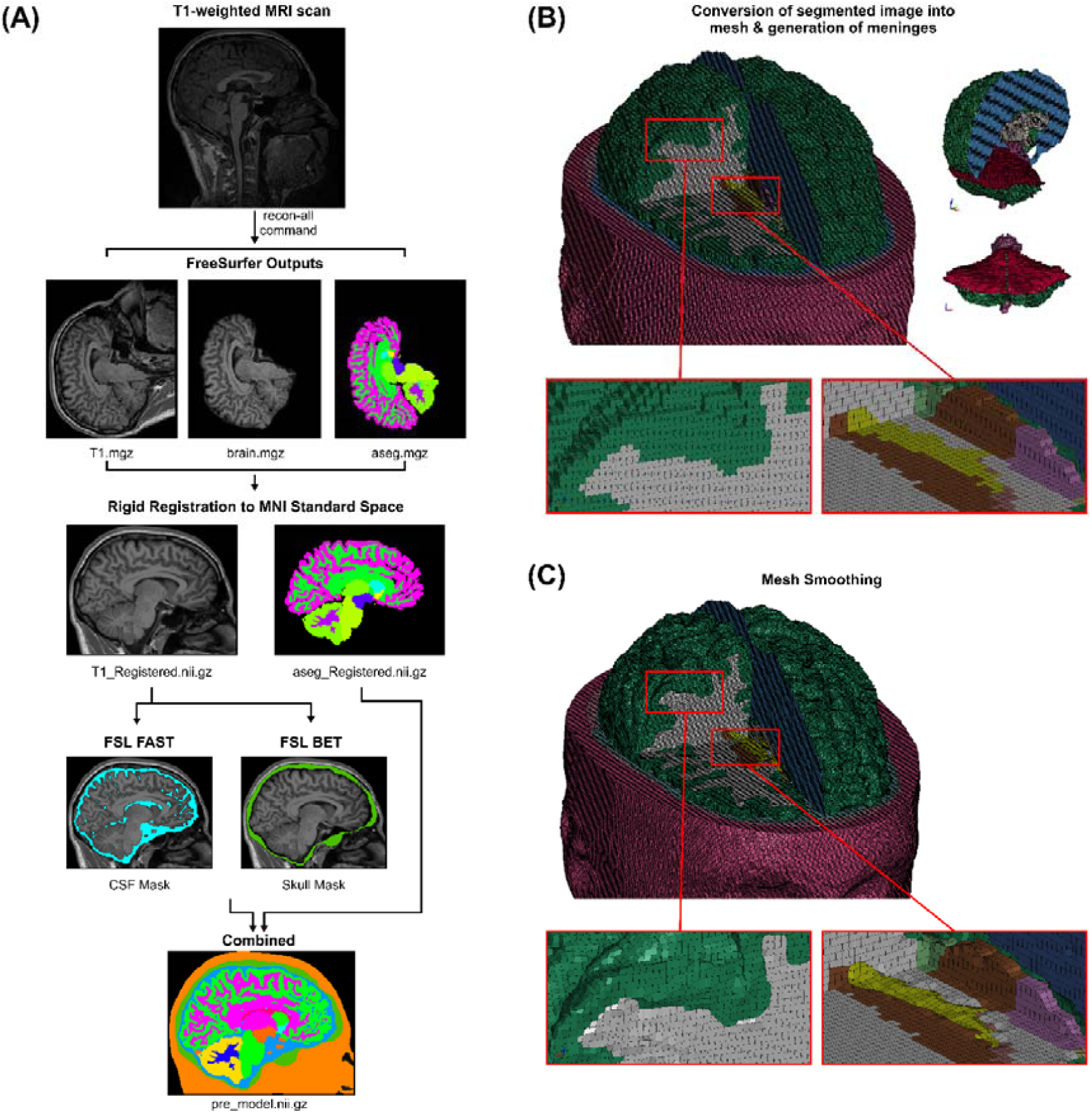
| Overview of the mesh generation pipeline. (**A**) Employing FreeSurfer’s recon-all command on a T1-weighted MRI scan provides a segmented brain parcellation (aseg.mgz) and a conformed T1-weighted image (T1.mgz). These outputs are then rigidly registered into MNI standard space. The registered T1 is then segmented again by FSL FAST and BET. The registered parcellation is combined with the FSL segmentation outputs to produce a comprehensive whole-head segmented image. **(B)** The final segmented image is directly converted into cubic hexahedral meshes, followed by the automatic generation of meninges, and **(C)** Cubic meshes were smoothed at the boundaries of regions to reduce stair-step effect.

### 2.2 Input to PARS: FreeSurfer’s recon-all outputs

The input to PARS is the outputs of FreeSurfer’s recon-all command applied to the T1-weighted MRI scan of the subject (Fischl, 2012). More specifically, three outputs are required by the pipeline: i) the conformed T1-weighted image (T1.mgz), ii) the skull-stripped brain volume (brain.mgz), and iii) the multi-label anatomical parcellation of cortical and subcortical structures (aseg.mgz) (Fig. 1A). FreeSurfer is freely available and well-documented, with full installation and usage instructions available at https://surfer.nmr.mgh.harvard.edu. The recon-all command, which typically requires 4 to 6 hours per subject on a standard CPU, is independent from the pipeline and can be performed in parallel across several subjects. The anatomical parcellation provides the detailed classification of brain structures, which forms the foundation for the finite element brain model.

### 2.3 Image pre-processing

Head orientation can be different in MRI scans, leading to head models that need to be re-oriented after creation. To avoid this, we first align the head with a standard MRI space. To achieve this, first PARS converts the FreeSurfer produced images to the standard NIfTI format, ensuring compatibility with FSL and other widely used neuroimaging tools (Fischl, 2012; Jenkinson et al., 2012). Then, it applies rigid registration (6 degrees of freedom) on the images, using FSL FLIRT (FMRIB’s Linear Image Registration Tool), to align them with the MNI152 standard space (Fig. 1A). Rigid registration applies translation and rotation only, thus avoiding any scaling or shearing of the image. This approach standardises the spatial orientation of the model while preserving the subject’s anatomical morphology.

### 2.4 Hybrid segmentation

Accurate segmentation is vital for defining the interfaces between distinct tissues, such as the ventricle-brain or brain-CSF-skull interfaces. To achieve this, we developed a hybrid segmentation strategy that integrates the strengths of two established neuroimaging libraries, FreeSurfer and FSL. FreeSurfer provides an accurate brain parcellation, but it cannot reliably segment CSF or the skull boundary. To overcome this limitation, PARS uses FSL FAST and FSL BET to accurately segment CSF, skull and skin and generate their masks (Fig. 1A).

Another limitation of FreeSurfer is that its brain mask can contain unclassified voxels (Fig. 2). The unclassified voxels are discontinuities in the segmentation that propagate into the FE mesh as void regions, acting as sites of unrealistic stress concentration during simulation. To resolve this issue, PARS performs a voxel-wise comparison between the two segmentation datasets generated by FreeSurfer and FSL and labels the “missing voxels”. Subsequently, to classify these unassigned voxels, we developed an iterative, neighbourhood-based algorithm consisting of two phases. First, the algorithm prioritises brain tissue by examining the neighbours of each missing voxel while explicitly ignoring CSF. If a specific brain tissue label accounts for more than 30% of neighbouring voxels, the missing voxel is assigned that identity. Second, a final cleanup step is applied to any remaining unassigned voxels; these are simply assigned the identity of their most frequent neighbour, which may now include CSF, ensuring there are no empty gaps in the volume. In parallel, the FSL-derived CSF mask is used to fill the subarachnoid space, producing a continuous fluid layer between the reconstructed brain surface and the inner skull.

**Fig. 2.**
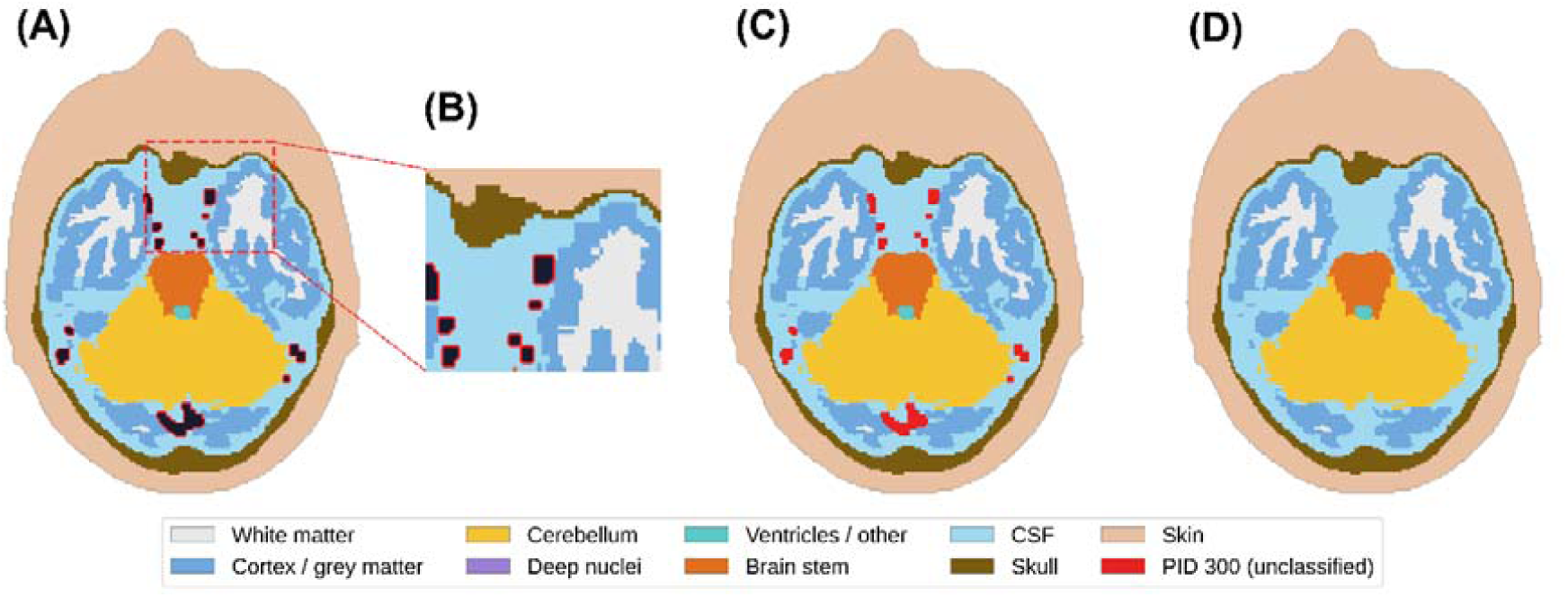
| Detection and reclassification of unclassified brain-tissue voxels. An axial slice (z = 49) from a subject, illustrating how PARS determines the voxels within the FSL brain mask which are not labelled in the FreeSurfer parcellation. **(A)** Combined segmentation prior to gap correction: The brain is segmented by FreeSurfer, while the surrounding CSF, skull, and skin regions are derived from FSL. The small black voids (outlined in red) represent voxels within the brain that were missed by FreeSurfer and remain unlabelled. The dashed box marks the region enlarged in (B). **(B)** Enlarged view showing the unclassified voids (black) surrounded by labelled tissue on all sides. **(C)** The same slice after the unclassified voxels are assigned the placeholder label PID 300 (red), making their spatial distribution explicit across the whole brain. **(D)** Final segmentation after iterative neighbourhood reclassification: each PID 300 voxel is reassigned to the most frequent tissue label in its 3×3×3 neighbourhood until no unclassified voxels remain, yielding a complete, gap-free parcellation ready for mesh generation.

### 2.5 Mesh Generation

PARS converts the segmented image into FE mesh using a direct voxel-to-element approach, generating a fully hexahedral mesh. Hexahedral elements, compared with tetrahedral elements used in some brain models, have superior performance in explicit simulations, providing better computational stability and resistance to volumetric locking in nearly incompressible materials such as brain tissue (Giudice et al., 2019; Tadepalli et al., 2011).

The element size of the generated mesh depends on the resolution of the MRI image, which varies across scanners and acquisition protocols. FreeSurfer’s default output images have a 1×1×1 mm³ resolution; the voxel-based FE mesh created from these images can have several million elements. While this fine resolution improves anatomical fidelity, the associated computational cost may be prohibitive for many users. Low resolution images, e.g. with voxel sizes of 1.5 mm or 2 mm, can be generated by resampling the high-resolution images. Creating models from these downsampled images can substantially reduce model size and simulation time, but with reduced anatomical accuracy. PARS allows users to specify the desired element size, which it uses to automatically resample the images and create FE meshes with the defined element size.

### 2.6 Algorithmic reconstruction of meningeal structures

FE brain meshes created using the voxel-to-element approach often omit meningeal structures due to their thin geometry indistinguishable at standard MRI resolutions (Chen and Ostoja-Starzewski, 2010). However, these structures are orders of magnitude stiffer than brain tissue (Betsholtz et al., 2024), and they play a significant role in brain biomechanics by limiting the relative motion of left and right hemispheres or constraining the upward movements of the corpus callosum (Darvishi et al., 2026; Hernandez et al., 2019). Although some studies have attempted to incorporate these structures manually in the brain model, these approaches compromise the automation (Miller et al., 2016). To address this issue, we developed an algorithm to determine the position of meninges based on anatomical landmarks and create them. This step is performed prior to the mesh smoothing process, because the algorithm can better identify anatomical landmarks using the clear orthogonal boundaries of the raw elements.

Falx: The algorithm identifies the longitudinal fissure separating left and right hemispheres. A shell layer is generated in the mid-sagittal plane, extending from the corpus callosum to the interior surface of skull.

Tentorium: The interface between the inferior surface of the occipital/temporal lobes and the superior surface of the cerebellum is identified and a shell layer is generated as a partition representing the tentorium.

Pia and Dura mater: The pia mater is generated as a shell layer coincident with the outer surface of the brain parenchyma, while the dura mater is generated from the inner surface of the skull mesh. Although the dura is rigidly tied to the skull, its inclusion lets the brain–skull interface be modelled as a shell-to-shell (dura–pia) rather than a solid-to-shell (skull–pia) contact, which can improve contact behaviour when simulating disorders like normal pressure hydrocephalus (Darvishi et al., 2026).

### 2.7 Mesh smoothing

A known limitation of direct voxel-to-element conversion is the creation of stair-step artifacts at interfaces. These artifacts, particularly at the brain/CSF interface, can lead to stress concentrations and numerical instabilities. To mitigate this, PARS applies a Laplacian smoothing algorithm adapted from the methodology presented in (Chen and Ostoja-Starzewski, 2010). The algorithm operates on the interface nodes of the generated mesh through an iterative process. The new position of a node *i* at iteration *n* + 1 is updated based on the position of its *N* neighbours

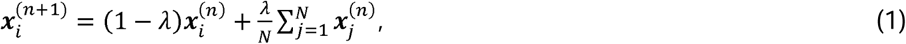

where *λ*, is the smoothing parameter. We recommend *λ*, = 0.3, consistent with (Chen and Ostoja-Starzewski, 2010), applied over 8 iterations.

Mesh smoothing, however, can adversely affect element quality and computational time. Node repositioning can distort elements, reducing simulation accuracy and causing simulation instability. Element distortion is quantified by the Jacobian ratio, which can be controlled during mesh smoothing. Additionally, node repositioning can reduce the characteristic length of elements, thereby decreasing the stable time step of explicit solvers, which are used in the vast majority of brain simulations, and increasing simulation wall-time. The stable time step (Δ*t*) is defined by the characteristic element length (*L_c_*) and speed of sound in the material (*c*):

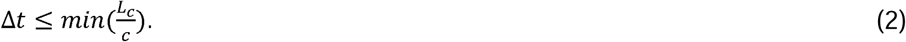

The speed of sound depends on element formulation. For 3D elements (such as skin, skull, and CSF), *c* is calculated as follows:

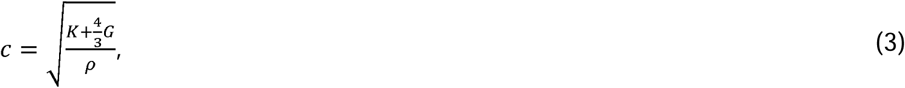

where *ρ* is density and K is bulk modulus. G is i) the shear modulus for linear elastic materials, ii) the small strain shear modulus for hyperelastic materials, and iii) the instantaneous shear modulus for viscoelastic materials. When a Prony series is used to describe viscoelastic behaviour (e.g., (Ghajari et al., 2017; Kleiven, 2007)), the instantaneous shear modulus is *G*_0_ = *G*_∞_ + *G*_1_ + *G*_2_ + … + *G_n_*, where *G_i_* are the Prony series amplitudes. For 2D elastic elements (such as falx, tentorium, dura mater, and pia mater), c is calculated as:

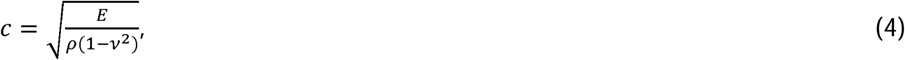

where *E*, is the elastic modulus and *v* is the Poisson’s ratio.

In order to control the element quality and stable time step during mesh smoothing, we incorporated a “node locking” algorithm within PARS. After each mesh smoothing iteration, the algorithm calculates the Jacobian ratio and stable time step of each element, and if they fall below a threshold, it “locks” the nodes of the elements, meaning that the position of its nodes cannot be moved further in following smoothing iterations. These thresholds are user-defined and directly determine the minimum mesh quality guaranteed after smoothing, allowing users to balance geometric conformity against element quality and stability according to their simulation requirements. A 0.2 threshold for the Jacobian ratio, 200 ns stable time step threshold for skin and skull elements, and a 2,000 ns stable time step threshold for all remaining elements are often suitable for simulation of brain biomechanical loading, e.g. in TBI and NPH. These different thresholds for skin/skull and other elements allow for increasing the simulation time in simulations where skin and skull are assumed to be rigid, e.g. when head kinematics is used for loading the head model in TBI.

### 2.8 Brain-skull contact repair

Direct voxel-to-element conversion can result in shared nodes between brain and skull elements, creating sites of non-physical brain-skull contact where no CSF layer is present (Fig. 3A and C). Under loading, these shared nodes produce sites of unrealistic strain in the brain parenchyma (Fig. 3B). To avoid such shared nodes, we developed a contact repair algorithm for PARS. It first iterates through the brain surface and detects any nodes shared between brain and skull. All skull elements associated with these shared nodes are then flagged and reclassified as CSF (Fig. 3C). This ensures that there is a continuous CSF layer between brain and skull.

**Fig. 3.**
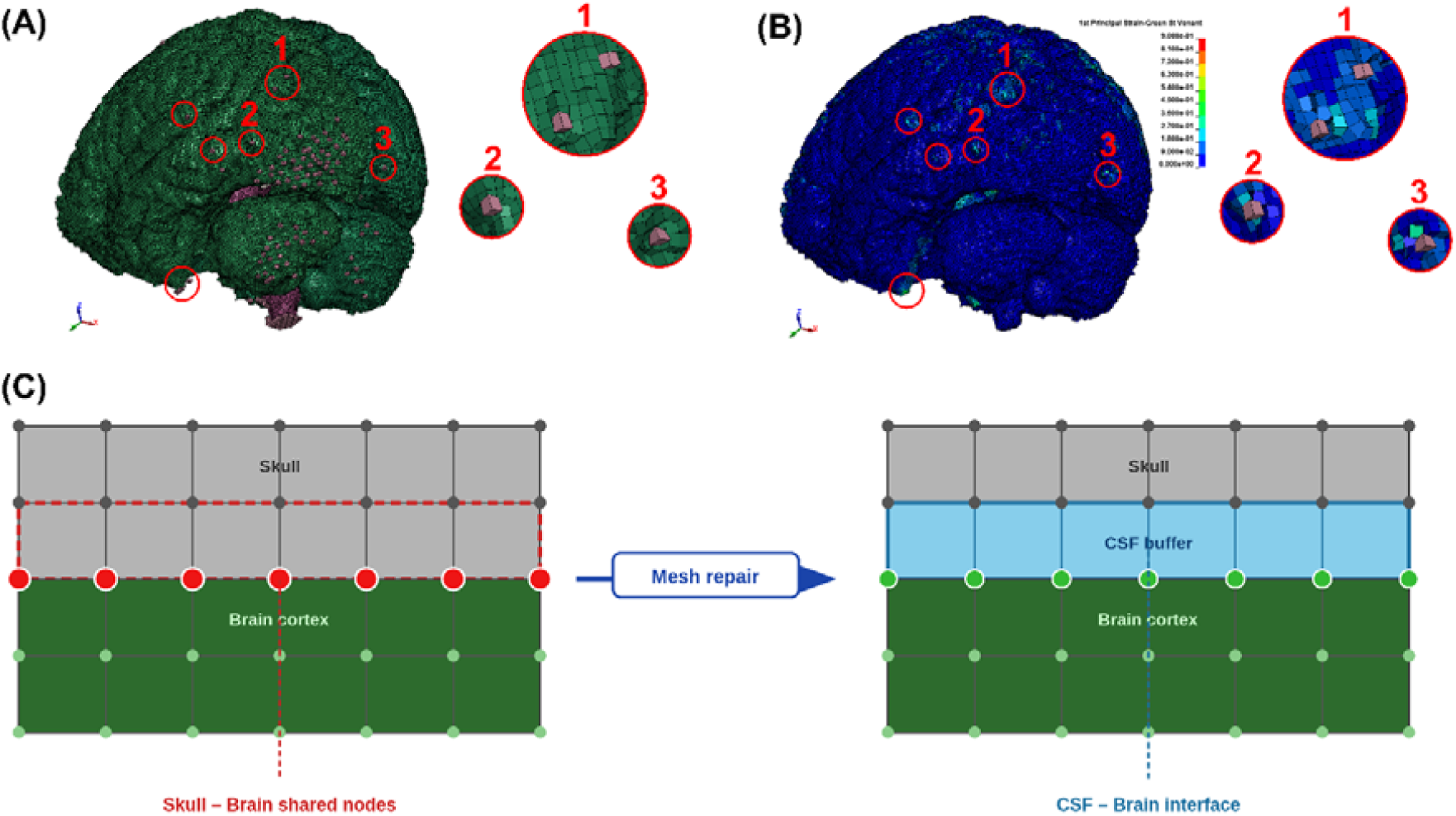
| Brain-skull contact repair in PARS. (**A**) Brain surface mesh following voxel-to-element conversion, with nodes shared between the brain and the skull, highlighted with red circles. **(B)** Maximum principal Green-Lagrange strain distribution in the brain under loading, demonstrating unrealistic strain concentrations localised at the sites of shared nodes. **(C)** Schematic of the mesh repair procedure. The mesh repair algorithm identifies such brain/skull shared nodes and relabels them as CSF elements.

### 2.9 Material assignment

The generated head mesh contains over 50 unique anatomical regions, distinguished via distinct part numbers. PARS currently assigns material properties to the distinct tissue classes where experimental data exists, including grey and white matter, CSF, ventricles, skull, skin, falx, tentorium, pia, and dura mater. Due to the lack of region-specific experimental data for deep brain sub-structures, these regions are currently assigned with the material properties of white matter. However, the distinct segmentation of these regions is preserved within the final mesh. This design supports future extension when novel experimental data for specific deep brain regions become available, allowing for the assignment of new material models and properties to these regions without the need for mesh regeneration.

### 2.10 Pipeline Evaluation and Robustness Testing

We evaluated the performance of PARS by creating head models for a diverse range of subjects. T1-weighted MRI scans from 23 subjects (9 females, age = 35.13 ± 13.80 years) with a wide range of cranial volumes (832.4 cm³ to 1,329 cm³) were used. Finite element meshes were generated for each subject at three target resolutions (1.0 mm, 1.5 mm, and 2.0 mm), with the material properties listed in Table 1 and Table 2 used to evaluate the dilatational wave speed and control minimum timestep during smoothing. There was no manual intervention throughout.

**Table 1.** Material properties of brain tissue.

| Tissue | Density [$\text{kg}/\text{m}^3$] | Shear Modulus $G$[MPa] | Bulk Modulus $K$[MPa] |
| --- | --- | --- | --- |
| Brain and brain stem | 1040 | 0.417097 | 50 |

**Table 2.** Material properties of the head tissues.

| Tissue | Density [$\text{kg}/\text{m}^3$] | Young's Modulus $E$[MPa] | Poisson's ratio |
| --- | --- | --- | --- |
| CSF (SAS and ventricles) | 1040 | 0.0015 | 0.49999 |
| Falx and tentorium | 1130 | 31.5 | 0.45 |
| Pia mater | 1130 | 11.5 | 0.45 |
| Skin | 1000 | 16.7 | 0.42 |
| Cortical bone | 1900 | 15000 | 0.21 |
| Trabecular bone | 1500 | 4600 | 0.05 |

Geometric fidelity was assessed by quantifying the relative volume difference between the smoothed hexahedral mesh and the segmented MRI for each anatomical regions across all three resolutions. To assess element quality, the scaled Jacobian ratio and aspect ratio were calculated for each hexahedral element across all subjects and resolutions, using the PyVista library in Python (Sullivan and Kaszynski, 2019). Explicit simulation feasibility was assessed by evaluating the stable timestep for all subjects and resolutions.

All computations were performed on a high-performance computing (HPC) cluster at Imperial College London. The Linux environment enabled the use of FreeSurfer and FSL command-line tools. Each job used only a single core of an AMD EPYC 7742 processor (64 cores, 2.25 GHz) and 8 GB of RAM. Given these minimal hardware requirements, PARS can be run efficiently on standard Linux workstations without the need for specialised computing infrastructure.

## 3 Results

### 3.1 Image pre-processing and hybrid segmentation for a female and a male subject

Pre-processing and segmentation outputs for one female and one male subject are shown in Fig. 4. The three images from FreeSurfer recon-all segmentation, shown in Fig. 4A, were the input to the pipeline. PARS re-oriented these brain volumes to align them with the MNI152 standard space (Fig. 4B). It then combined the registered FreeSurfer parcellation with FSL FAST and BET hybrid segmentation outputs to produce a complete segmented image representing all intracranial compartments, including grey matter, white matter, CSF, ventricles, skull, and skin (Fig. 4C). When tested on all 23 subjects, the pre-processing and segmentation steps required approximately 5 minutes per subject.

**Fig. 4.**
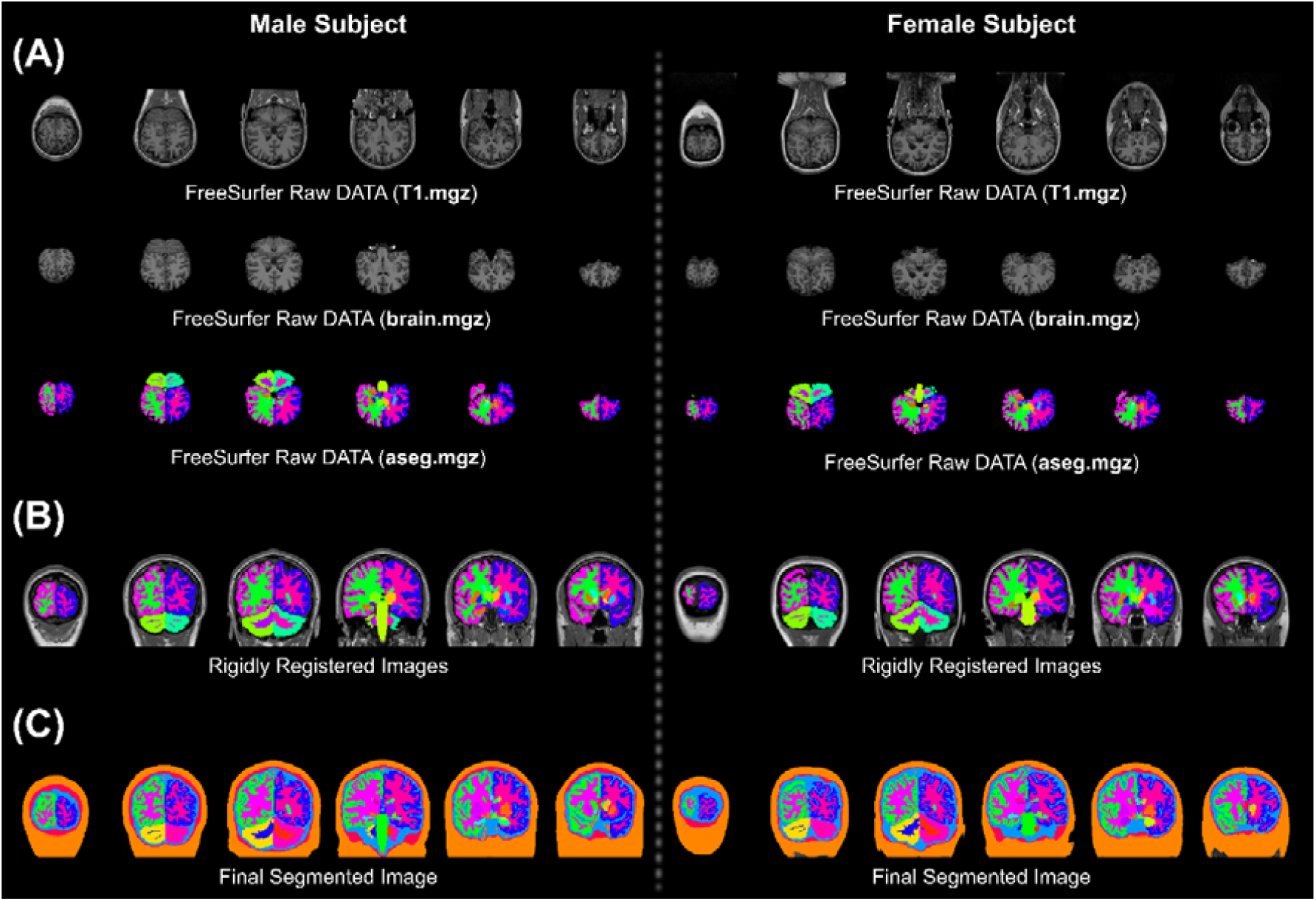
| Image pre-processing and hybrid segmentation. (**A**) Raw data extracted from recon-all FreeSurfer segmentation, **(B)** FreeSurfer data rigidly registered into MNI152 space to standardise the orientation and **(C)** Combination of FSL’s FAST and BET segmentations with registered FreeSurfer segmentation to create the final head segmentation.

### 3.2 Automatic mesh generation and meningeal reconstruction

PARS generated hexahedral meshes using the segmented images and successfully reconstructed the meningeal structures, including the falx, tentorium, pia mater, and dura mater (Fig. 5). As shown in Fig. 5A, the generated falx was closely aligned with the longitudinal fissure, while the tentorium correctly partitioned the cerebrum from the cerebellum along the transverse fissure. The created pia and dura mater are also shown in Fig. 5B and Fig. 5C.

**Fig. 5.**
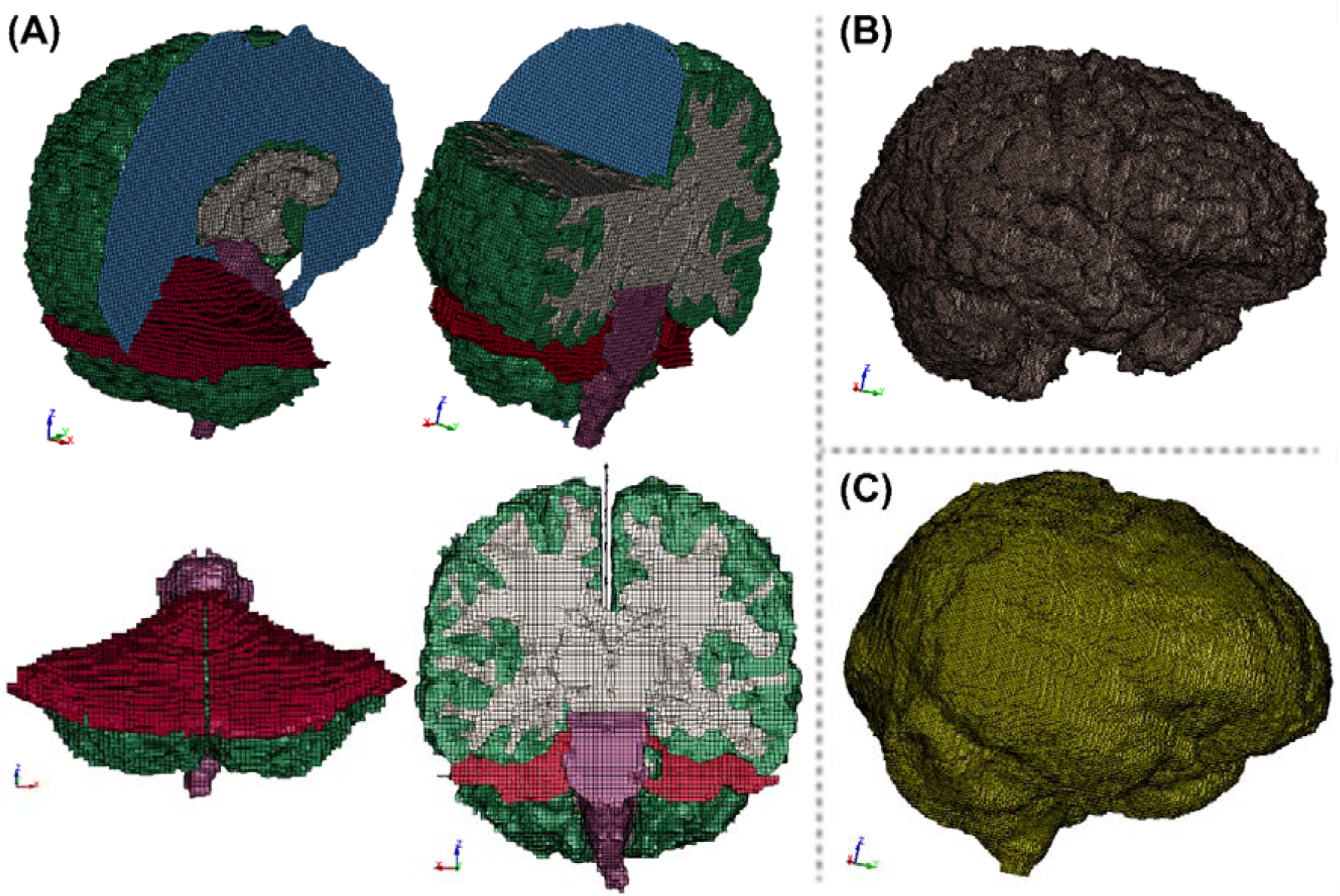
| Meningeal geometries generated automatically by the pipeline. (**A**) Anatomy of the falx (blue) and tentorium (red) and how they separate left and right hemispheres and cerebrum from the cerebellum, respectively, **(B)** Anatomy of pia mater represented as shell elements and **(C)** Anatomy of dura mater represented as shell elements.

### 3.3 Mesh smoothing

The result of the Laplacian smoothing is shown in Fig. 1C. For this step, PARS used the following parameters: λ = 0.3, number of iterations = 8, the Jacobian ratio limit = 0.2, skin/skull stable timestep limit = 200 ns, and brain stable timestep limit = 2000 ns. The figure shows that PARS successfully smoothed the mesh at different interfaces, such as skull-CSF, CSF-brain and brain-ventricles.

### 3.4 Mesh Quality

Mesh quality, measured by the Scaled Jacobian (SJ) and Aspect Ratio (AR), was generally high across all subjects and resolutions, as shown in Fig. 6A. At 1 mm resolution, the median Scaled Jacobian was 0.976 ± 0.012, with 88.6 ± 0.9% of elements having SJ ≥ 0.5. The median Scaled Jacobian decreased modestly to 0.933 ± 0.018 for 1.5 mm resolution and 0.921 ± 0.016 for 2 mm resolution, but the percentage of elements with SJ ≥ 0.5 increased slightly. Similarly, the aspect ratio across all subjects was slightly better (closer to 1) for the 1 mm resolution (1.225 ± 0.077) than 1.5 mm (1.380 ± 0.046) and 2 mm (1.400 ± 0.034), but the percentage of elements with AR < 3 was slightly higher for 1.5 mm and 2 mm resolutions.

**Fig. 6.**
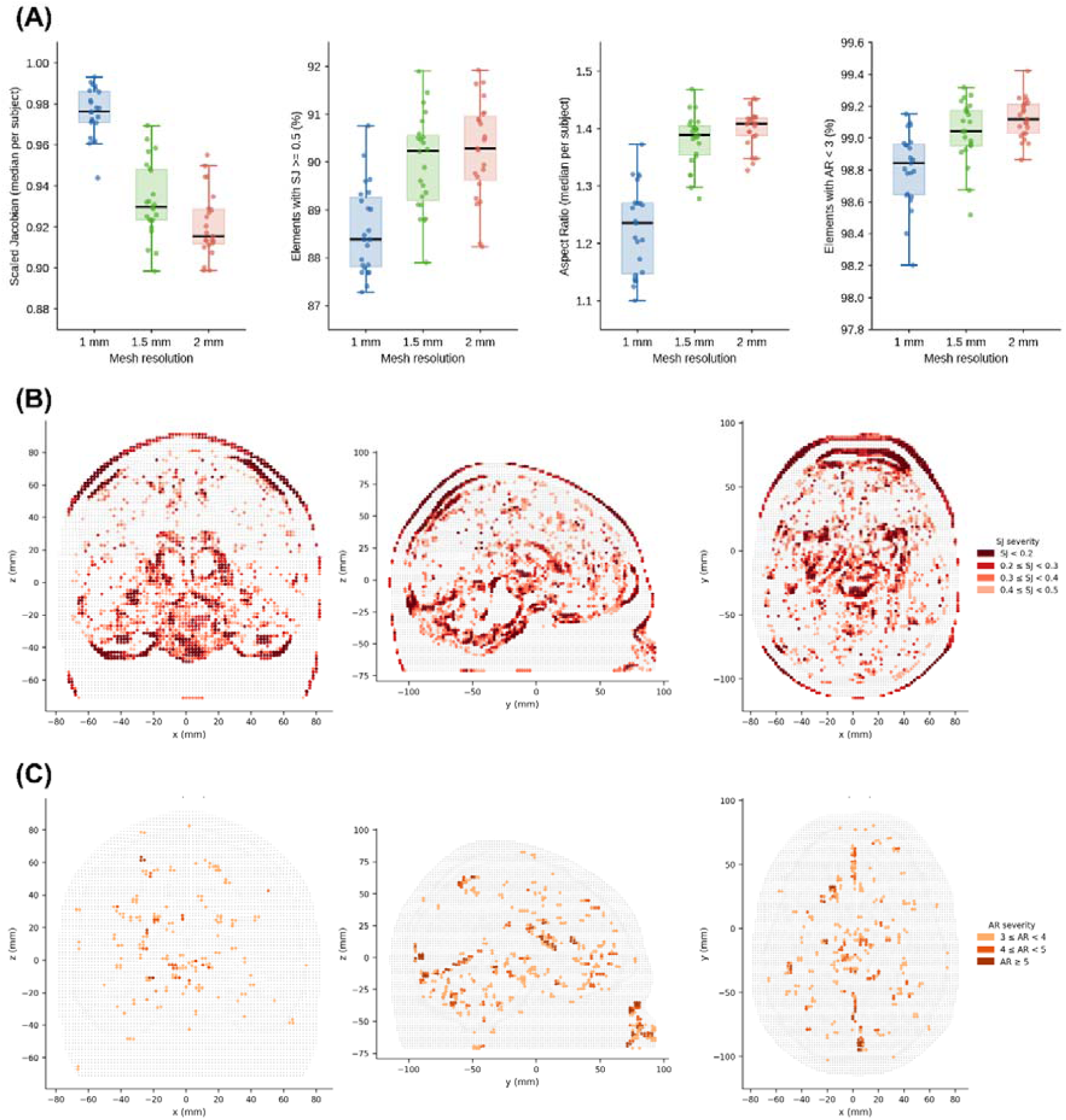
| Mesh quality assessment across 23 subjects at three mesh resolutions. (**A**) Per-subject values (dots) overlaid on group box plots (median, IQR, and range). From left to right: Scaled Jacobian (SJ) median; proportion of elements with SJ ≥ 0.5; Aspect Ratio (AR) median; proportion of elements with AR < 3. Higher SJ and lower AR indicate better element conditioning; finer meshes (1 mm) consistently yield higher SJ medians. **(B)** Elements with SJ < 0.5 (8.1% of total) and **(C)** Elements with AR ≥ 3 (0.6% of total), for a representative subject (2 mm; 427,905 elements), shown as ±8 mm slabs through the coronal, sagittal, and axial mid-planes. Grey: all elements in the slab. Low-quality elements concentrate at the skull–CSF–brain interfaces and ventricular surfaces.

The spatial distribution of lower-quality elements is shown in Fig. 6B for a representative subject at 2 mm resolution. Elements with SJ < 0.5 (8.1% of elements in this subject) and those with AR ≥ 3 (0.6%) were concentrated at the skull–CSF–brain interfaces and ventricular surfaces.

### 3.5 Stable timestep

The stable timestep for the 1 mm, unsmoothed skull/skin was 348 ns (Fig. 7A, smoothing iteration = 0) and for the unsmoothed brain/CSF was 4535 ns (Fig. 7B, smoothing iteration = 0). With every smoothing iteration, the stable timestep reduced but with varying rates for different mesh resolutions. Fig. 7 also shows that when the stable timestep fell below the threshold defined for locking nodes, it remained nearly constant, showing that the node locking was triggered and effectively limited further reduction in the stable timestep.

**Fig. 7.**
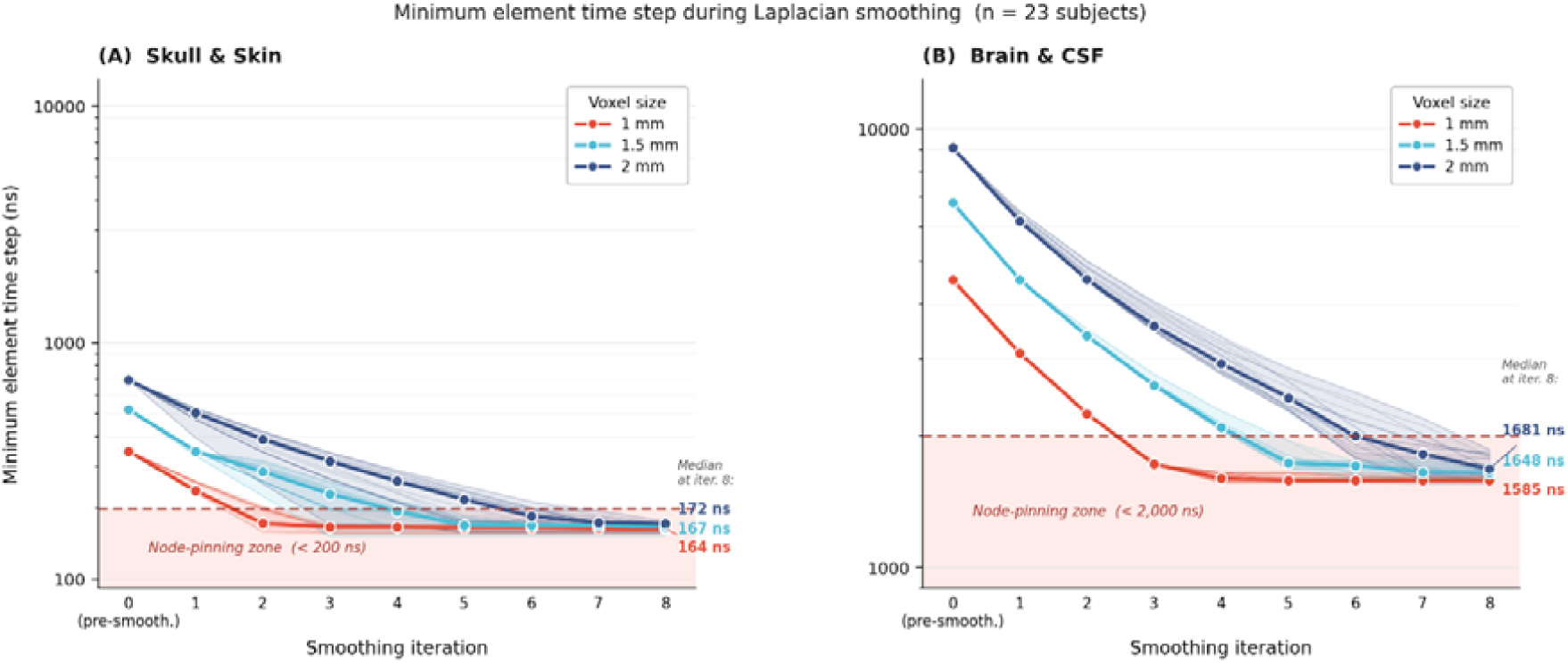
| Minimum element time step during Laplacian smoothing (n = 23 subjects at three mesh resolutions). (**A**) Skull and skin elements: per-subject trajectories (faint lines) and median across subjects (bold line with circles) at each smoothing iteration. The shaded region indicates the node-pinning zone (< 200 ns). Median time steps at iteration 8 were 164 ns, 167 ns, and 172 ns at 1 mm, 1.5 mm, and 2 mm resolutions, respectively. (B) Brain and CSF elements: per-subject trajectories (faint lines) and median across subjects (bold line with circles) at each smoothing iteration. The shaded region indicates the node-pinning zone (< 2,000 ns). Median time steps at iteration 8 were 1,585 ns, 1,648 ns, and 1,681 ns at 1 mm, 1.5 mm, and 2 mm resolutions, respectively.

### 3.6 Volumetric accuracy across mesh resolutions

At 1 mm resolution, the relative volume difference between the smoothed mesh and segmented image remained within ±5% for all tissue compartments, with the largest deviation observed as a volume increase in the brain stem (+5.07 ± 1.26%), as shown in Fig. 8. At coarser resolutions, white matter and CSF showed the greatest sensitivity: white matter volume decreased to −12.97 ± 2.29% at 2 mm, while CSF volume increased, reaching a difference of +16.34 ± 2.74%.

**Fig. 8.**
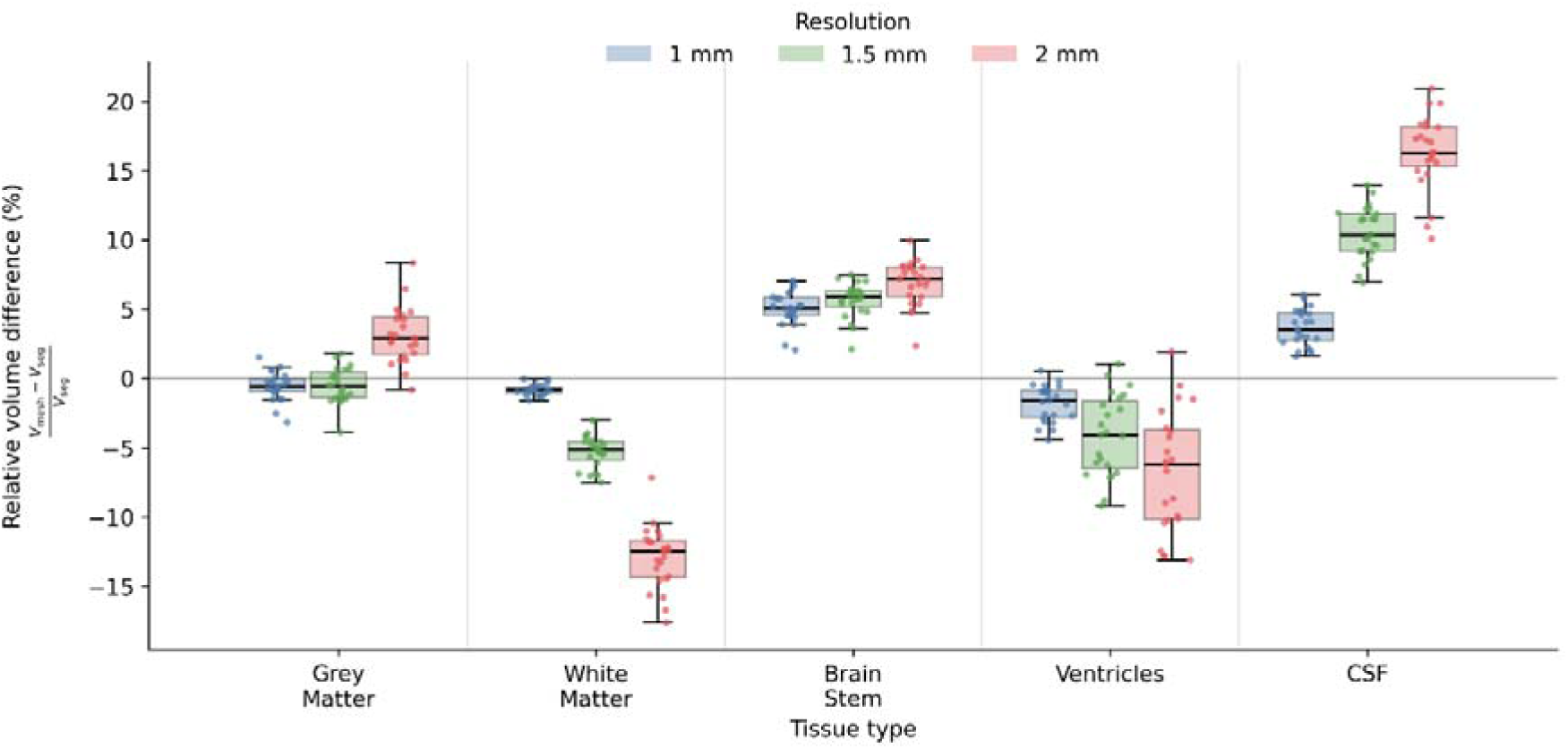
| Relative volume error (%) between the generated hexahedral mesh and the segmented MRI, shown per tissue type across three mesh resolutions. Each group shows per-subject values (dots) overlaid on group statistics for 1 mm (blue), 1.5 mm (green), and 2 mm (red) resolutions. The horizontal line at 0% indicates perfect volumetric correspondence.

### 3.7 Runtime of PARS’ different stages

The total pipeline runtime, from the point of reading the FreeSurfer output files to the generation of smoothed head mesh, ranged between approximately 9 minutes and 38 minutes depending on the mesh resolution (Table 3). Mesh generation, which includes reconstruction of meninges, was the most time-consuming stage, accounting for approximately 81%, 61%, and 41% of the total runtime at 1 mm, 1.5 mm, and 2 mm resolutions, respectively. At 2 mm resolution, image pre-processing and segmentation became the dominant stage, reflecting the reduced mesh generation cost at coarser resolutions.

**Table 3.**
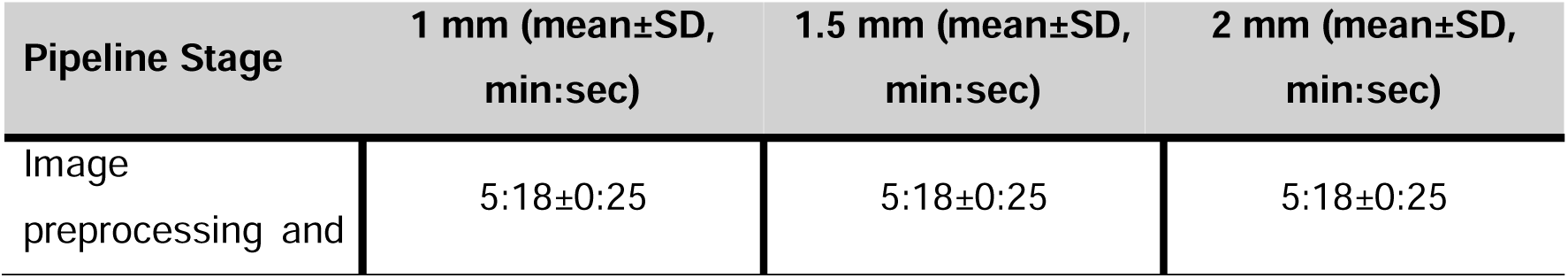
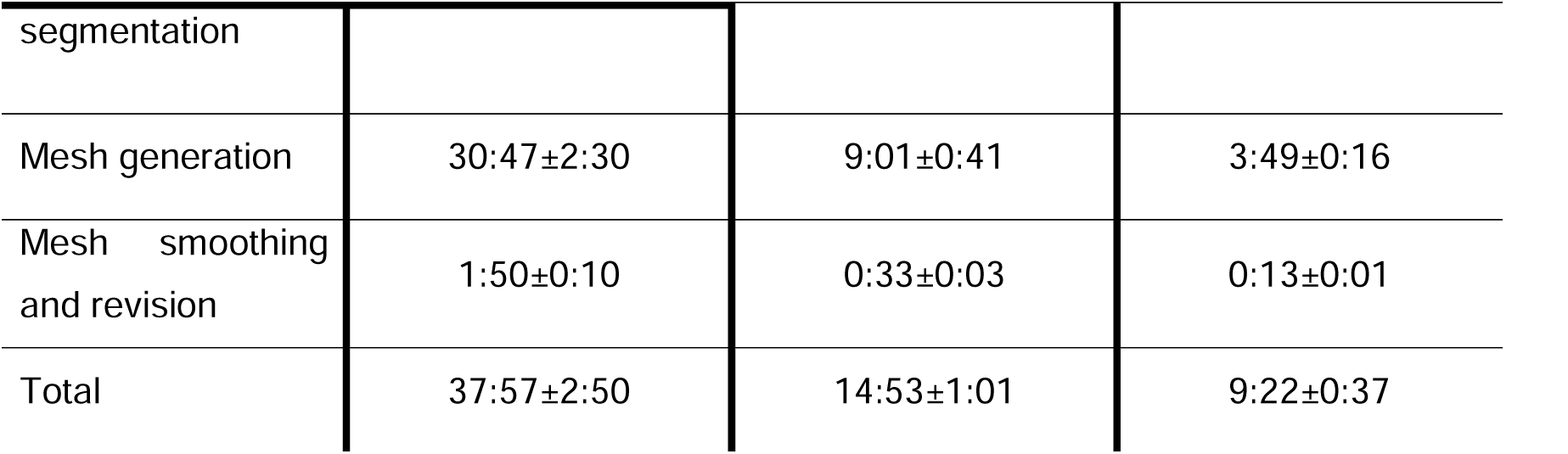
Time required for different steps of PARS.

## 4 Discussion

We have developed an automated and open-source pipeline that enables the generation of anatomically detailed, subject-specific head models directly from standard T1-weighted MRI images. Unlike previous model creation pipelines that require manual intervention or geometry simplification, PARS preserves complex anatomical details without any user input. Its successful application to 23 subjects spanning a diverse range of cranial volumes, including automated reconstruction of falx and tentorium, demonstrates the pipeline’s robustness across a demographically diverse cohort. By releasing PARS as a freely available, open-source resource, we aim to lower the technical and financial barriers to subject-specific brain modelling and facilitate transparent, reproducible research into the biophysics of neurological conditions.

The models generated by the PARS pipeline have been validated in previous studies (Darvishi et al., 2026; Ghajari et al., 2017; Zimmerman et al., 2023). The models’ predictions of brain displacement have been compared against cadaveric experiments, achieving an average CORA score of 0.60 across multiple rapid head rotations (Alshareef et al., 2020, 2018; Duckworth et al., 2022). In addition, these models have been utilised in predicting the location of strain and strain rate concentration produced by head impacts, which overlapped with the location of neurodegenerative disease, CTE, pathology and microbleeds (Duckworth et al., 2022; Ghajari et al., 2017). These models have also predicted disproportionately large strain rates in brainstem regions involved in arousal for players who lost consciousness, providing a biomechanical explanation for loss of consciousness (Zimmerman et al., 2023). The clinical utility of the models generated by PARS has further been supported by their success in simulating the complex morphological changes associated with iNPH. Specifically, the model accurately predicted diagnostic biomarkers of iNPH and patterns of white matter abnormalities observed in MRI images of iNPH patients (Darvishi et al., 2026; Del Giovane et al., 2026). Together, these studies confirm the reliability of the PARS generated head models for studying a wide range of neurological conditions.

Open accessibility is a defining design principle of PARS. While several head models have been developed in the past decade, their codebases often remain inaccessible or depend on expensive commercial meshing software. By developing codes for executing hybrid segmentation and mesh creation and using established, freely available neuroimaging suites like FSL and FreeSurfer, we eliminate these barriers. Open-source availability also enables independent validation and replication by research groups, which is essential for rigorous biomechanical science. Ultimately, a zero-cost workflow creates a more inclusive research environment and supports the translation of subject-specific modelling from an academic tool to a clinical tool.

Another distinguishing feature of PARS is its high level of automation in the generation of not only the brain parenchyma, but also the meningeal structures, CSF, skull, and skin. In many existing model creation workflows (Chen and Ostoja-Starzewski, 2010; Li et al., 2021; Miller et al., 2016), meningeal structures either remain absent or require manual creation, which limits the ability to generate several models for large-scale cohorts. Given that these meningeal tissues significantly affect the biomechanical response of the brain to loading, their automated inclusion is vital for biofidelic simulations of conditions, such as TBI and iNPH. The successful generation of these tissues for all 23 subjects confirms the robustness of the anatomical feature detection algorithm used in PARS, demonstrating its ability to generate meningeal structures without manual intervention.

This automation is complemented by the anatomical fidelity of the generated models. The hybrid segmentation strategy, combining FreeSurfer’s anatomical parcellation with FSL’s voxel-wise probabilistic tissue maps, enables the incorporation of detailed anatomy of different compartments in the head model. CSF, as an example, is produced by integrating voxel-wise probabilistic maps from FSL FAST, rather than being approximated as a uniform layer of constant thickness, a simplification adopted in conventional models that fail to capture the non-uniform fluid distribution within the sulci (Miller et al., 2016). Some anatomical details play a key role in predicting deformation patterns. For instance, in iNPH, where ventricular enlargement pushes the brain tissue against the skull, the nonuniform distribution of CSF around the brain has allowed for predicting brain deformation and strain patterns that potentially explain brain morphology and white matter abnormality seen in iNPH patients (Darvishi et al., 2026). As another example, incorporating sulci in the model has enabled the prediction of large strains in locations of the chronic traumatic encephalopathy pathology (Ghajari et al., 2017; Zimmerman et al., 2023).

While PARS is fully automated, it benefits from a modular architecture that permits user intervention for anatomical refinement. Automated segmentation suites, including FreeSurfer, occasionally fail to resolve fine-scale structures. An example is the septum pellucidum, a small anatomical region where pathology has been observed in TBI and iNPH (Darvishi et al., 2026; Kamps et al., 2025), but often not segmented by FreeSurfer. Our pipeline addresses this issue by allowing users to verify and revise segmentation masks prior to mesh generation. Fig. 9 shows that by revising the brain mask, we were able to incorporate septum pellucidum in the brain mesh. This capability is one of the key differences between PARS and template-based morphing methods. Morphing techniques can rapidly transform a base model mesh into an individual’s space (Giudice et al., 2020; Ji et al., 2011; Li et al., 2021). However, these models are inherently dependent on the anatomical fidelity of the base model. If an anatomical feature, such as the septum pellucidum, is absent from the base model, that omission is propagated to all subject-specific models. In contrast, by allowing for the manual revision of segmentation masks, PARS can incorporate fine anatomical regions or abnormal regions, such as tumours, in the model.

**Fig. 9.**
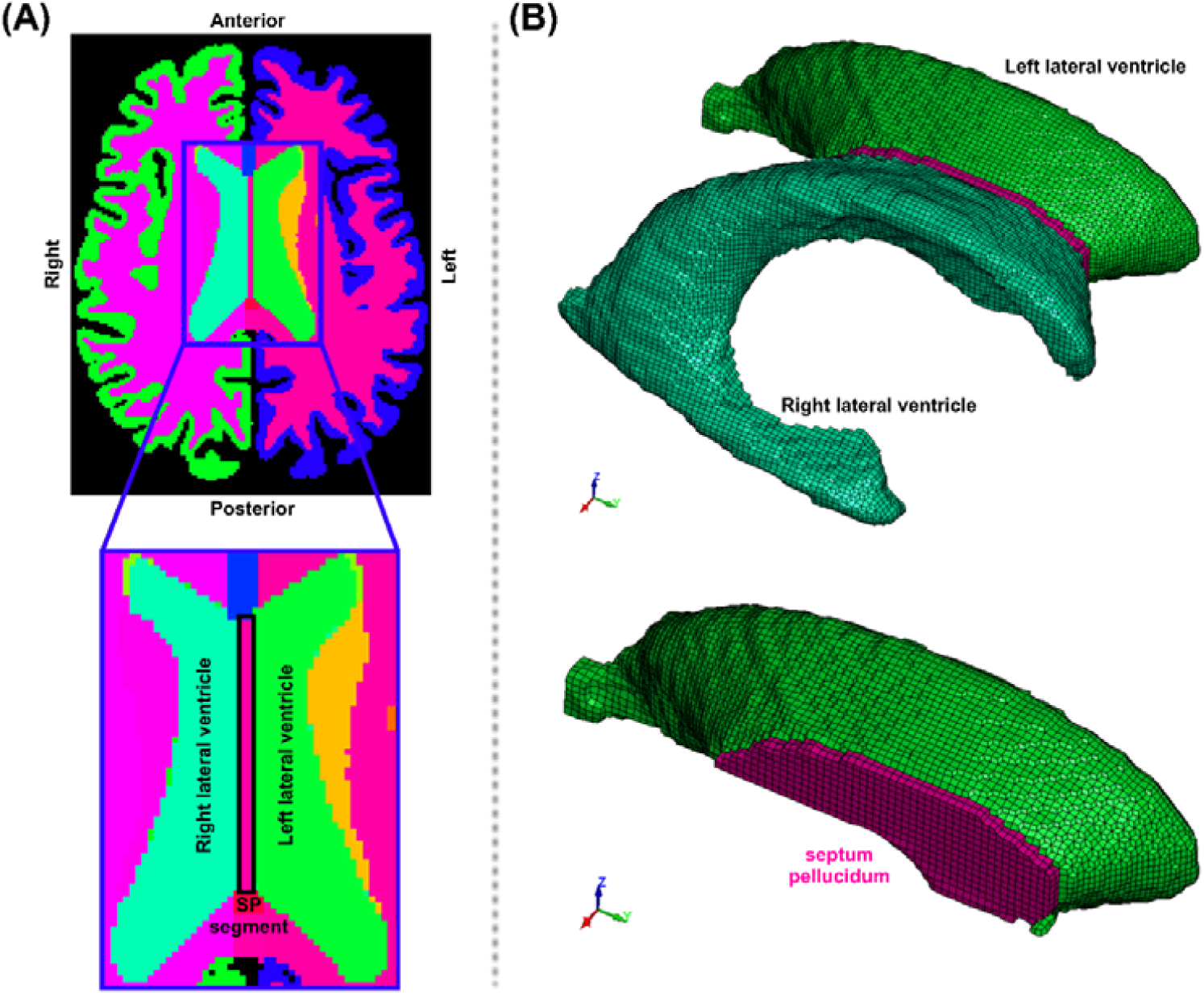
| Segmentation refinement for anatomical enhancement of the brain model. (**A**) Refining segmentation of septum pellucidum and **(B)** Anatomy of septum pellucidum after refinement, the region that has been omitted in previous FE models of the brain.

PARS provides direct control over mesh density and smoothing parameters, while controlling element quality and stable timestep. This allows researchers to control the trade-off between mesh resolution, mesh quality and computational runtime. This was enabled by locking the position of the nodes of elements that passed a threshold value for the Jacobian ratio and stable timestep, as shown for all 23 head models created. Having control over the stable time step does not exist in previous pipelines and commercial software (HyperWorks, n.d.); while they can smooth the mesh, they cannot control the stable timestep, leading to models that require unnecessarily long computational runtime. Our results confirm that node-locking was active during smoothing and that the user-defined thresholds provided effective, predictable control over the extent of mesh deformation.

The total intracranial volume of the PARS-generated meshes differed from the segmentation by only 0.54 ± 0.19% at 1 mm (rising to 1.15 ± 0.46% at 2 mm). This agreement is comparable to values reported for other validated subject-specific FE brain models (Giudice et al., 2021). In addition to total intracranial volume, we report per-tissue volume differences, a more detailed assessment than is typically provided. At 1 mm resolution, all anatomical regions were within approximately 5% (the largest deviation being the brain stem). At coarser resolutions, CSF showed the largest volume difference, which can partly be attributed to the brain-skull contact repair, which reclassifies skull elements as CSF at sites of direct brain-skull contact (Fig. 3), locally increasing the CSF volume in the mesh.

This work has some limitations. PARS requires segmented images produced by FreeSurfer, which typically requires 4 to 6 hours to fully segment a T1 image. Novel image segmentation approaches based on deep learning provide promising opportunities for substantially reducing this delay and improving the segmentation accuracy, which is propagated to the head mesh (Billot et al., 2023). Another limitation is the accuracy of the skull model. While the automated pipeline generates skull boundaries for predicting intracranial responses, the skull mesh may not be suitable for predicting skull fractures. If predicting skull fracture is the aim, more accurate models of the skull need to be generated by using other imaging modalities, such as CT. Recent studies have demonstrated the feasibility of generating high-fidelity skull models capable of predicting fracture with good accuracy (Henningsen et al., 2024; Lindgren et al., 2023). A limitation of the meningeal reconstruction algorithm is that the generated falx cerebri follows a planar approximation and cannot conform to a curved interhemispheric fissure. In subjects where significant curvature is present, this may result in the falx mesh intersecting adjacent brain tissue. Developing a surface-fitted meningeal generation approach that follows subject-specific fissure geometry is therefore an important direction for future work. Another limitation is that the pipeline cannot determine the head’s centre of mass, which is needed for simulations where the head is loaded by applying head kinematics to its centre of mass. Finally, while the pipeline was applied to 23 subjects spanning a range of cranial volumes and demographic profiles, evaluation on a larger and more diverse cohort would further strengthen confidence in its robustness and generalisability.

## 5 Conclusions

PARS addresses a longstanding barrier in computational biophysics: the absence of a fully automated, open-source pathway from MRI scan to simulation-ready FE head model. By combining hybrid segmentation, voxel-based meshing, algorithmic meningeal reconstruction, and user-controlled smoothing within a single pipeline built entirely on freely available tools, PARS makes subject-specific brain modelling accessible to researchers regardless of their institutional resources or software budget. Applied to 23 subjects without any manual intervention, the pipeline produced meshes of high geometric fidelity. Models generated using this framework have been validated against cadaveric experiments and demonstrated clinical utility in TBI and iNPH research. PARS holds promise to increase the use of subject-specific brain modelling across a range of neurological applications, e.g. TBI prediction, neurosurgery planning, and drug delivery optimisation, thus advancing mechanistic understanding of these conditions and ultimately their clinical care.

## 6 Acknowledgements

This work has been developed over several years with support from multiple funding sources. V.D. acknowledges the Design Engineering PhD Scholarship, Imperial College London. M.G. acknowledges the support of the Royal Academy of Engineering under the Senior Research Fellowship scheme (RCSRF2324-17-19). D.J.S. has received funding from the Medical Research Council, National Institute of Health Research, Alzheimer’s Society, Football Association, Rugby Football Union, and Premier League Rugby. H. D. acknowledges support from the Engineering and Physical Sciences Research Council (EPSRC) [EP/N509486/1 and reference 2024686]. E.Y.K.C. acknowledges postgraduate funding supplied by Sports and Wellbeing Analytics Ltd. and support from the Research Computing and Data Science Exemplars (ReCoDE) programme, funded by the Early Career Researcher Institute at Imperial College London. T.D.P. has been supported by a NIHR Clinical Lectureship, the Medical Research Council (MRC) (TBI-TS –UKRI2548), and by a UK Dementia Trials Network fellowship. We also thank the Imperial College Research Computing Service for computational resources (DOI: 10.14469/hpc/2232).

## 7 Ethics and inclusion statement

The acquisition of MRI data for this study was approved by the Health Research Authority’s London-Surrey Borders Research Ethics Committee (19/LO/0102), the London-Central Research Ethics Committee (18/LO/0249), the Camberwell St Giles Research Ethics Committee (17/LO/2066), and the Imperial College Research Ethics Committee (17IC4009). All participants provided digital or written informed consent.

## 8 Data Availability

All relevant data supporting the results in this study are available from the corresponding author upon reasonable request and following the completion of a material transfer agreement with Imperial College London.

## 9 Code Availability

The brain mesh generation pipeline used in this study (PARS) is openly available as a documented exemplar, including the complete MRI-to-mesh workflow and Jupyter notebooks, at https://headlabic.github.io/PARS/.

## 10 CRediT authorship contribution statement

**Vahid Darvishi**: Conceptualization, Methodology, Software, Formal analysis, Investigation, Data curation, Visualization, Validation, Writing – original draft, Writing – review & editing. **Emily Yik Kwan Chan**: Data curation, Validation, Writing – review & editing. **Harry Duckworth**: Software, Writing – review & editing. **Thomas D. Parker:** Data curation, Investigation, Writing – review & editing. **David J. Sharp**: Resources, Supervision, Funding acquisition, Writing – review & editing. **Mazdak Ghajari**: Conceptualization, Methodology, Software, Supervision, Funding acquisition, Project administration, Writing – review & editing.

## 11 Declaration of competing interests

D.J.S. serves on the Alzheimer’s Society Research Advisory Board and the Rugby Football Union Concussion Advisory Board. D.J.S. serves on the Editorial Board of the journal Brain. D.J.S. has received research contributions from Alamar, although unrelated to this work. The other authors declare that they have no known competing financial interests or personal relationships that could have appeared to influence the work reported in this paper.

## FIG

## Notes

https://headlabic.github.io/PARS/

### Competing Interest Statement

D.J.S. serves on the Alzheimer Society Research Advisory Board and the Rugby Football Union Concussion Advisory Board. D.J.S. serves on the Editorial Board of the journal Brain. D.J.S. has received research contributions from Alamar, although unrelated to this work. The other authors declare that they have no known competing financial interests or personal relationships that could have appeared to influence the work reported in this paper.

